# Genomic diversity of *Escherichia coli* isolates from non-human primates in the Gambia

**DOI:** 10.1101/2020.02.29.971309

**Authors:** Ebenezer Foster-Nyarko, Nabil-Fareed Alikhan, Anuradha Ravi, Gaëtan Thilliez, Nicholas Thomson, David Baker, Gemma Kay, Jennifer D. Cramer, Justin O’Grady, Martin Antonio, Mark J. Pallen

## Abstract

Increasing contact between humans and non-human primates provides an opportunity for the transfer of potential pathogens or antimicrobial resistance between host species. We have investigated genomic diversity, and antimicrobial resistance in *Escherichia coli* isolates from four species of non-human primate in the Gambia: *Papio papio* (n=22), *Chlorocebus sabaeus* (n=14), *Piliocolobus badius* (n=6) and *Erythrocebus patas* (n=1). We performed Illumina whole-genome sequencing on 101 isolates from 43 stools, followed by nanopore long-read sequencing on eleven isolates. We identified 43 sequence types (STs) by the Achtman scheme (ten of which are novel), spanning five of the eight known phylogroups of *E. coli*. The majority of simian isolates belong to phylogroup B2—characterised by strains that cause human extraintestinal infections—and encode factors associated with extraintestinal disease. A subset of the B2 strains (ST73, ST681 and ST127) carry the *pks* genomic island, which encodes colibactin, a genotoxin associated with colorectal cancer. We found little antimicrobial resistance and only one example of multi-drug resistance among the simian isolates. Hierarchical clustering showed that simian isolates from ST442 and ST349 are closely related to isolates recovered from human clinical cases (differences in 50 and seven alleles respectively), suggesting recent exchange between the two host species. Conversely, simian isolates from ST73, ST681 and ST127 were distinct from human isolates, while five simian isolates belong to unique core-genome ST complexes—indicating novel diversity specific to the primate niche. Our results are of public health importance, considering the increasing contact between humans and wild non-human primates.

**Impact statement:** Little is known about the population structure, virulence potential and the burden of antimicrobial resistance among *Escherichia coli* from wild non-human primates, despite increased exposure to humans through the fragmentation of natural habitats. Previous studies, primarily involving captive animals, have highlighted the potential for bacterial exchange between non-human primates and humans living nearby, including strains associated with intestinal pathology. Using multiple-colony sampling and whole-genome sequencing, we investigated the strain distribution and population structure of *E. coli* from wild non-human primates from the Gambia. Our results indicate that these monkeys harbour strains that can cause extraintestinal infections in humans. We document the transmission of virulent *E. coli* strains between monkeys of the same species sharing a common habitat and evidence of recent interaction between strains from humans and wild non-human primates. Also, we present complete genome assemblies for five novel sequence types of *E. coli*.

**Author notes:** All supporting data, code and protocols have been provided within the article or through supplementary data files. Nine supplementary figures and six supplementary files are available with the online version of this article.

**Abbreviations:** ExPEC, Extraintestinal pathogenic *Escherichia coli*; ST, Sequence type; AMR, Antimicrobial resistance; MLST, Multi-locus sequence typing; VFDB, Virulence factors database; SNP, single nucleotide polymorphism; SPRI, Solid phase reversible immobilisation.

**Data summary:** The raw sequences and polished assemblies from this study are available in the National Center for Biotechnology Information (NCBI) Short Read Archive, under the BioProject accession number PRJNA604701. The full list and characteristics of these strains and other reference strains used in the analyses are presented in Table 1 and Supplementary Files 1-4 (available with the online version of this article).

## Introduction

*Escherichia coli* is a highly versatile species, capable of adapting to a wide range of ecological niches and colonising a diverse range of hosts (1, 2). In humans, *E. coli* colonises the gastrointestinal tract as a commensal, as well as causing intestinal and extraintestinal infection (2). *E. coli* is also capable of colonising the gut in non-human primates (3–5), where data from captive animals suggest that gut isolates are dominated by phylogroups B1 and A, which, in humans, encompass commensals as well as strains associated with intestinal pathology (6–9). *E. coli* strains encoding colibactin, or cytotoxic necrotising factor 1 have been isolated from healthy laboratory rhesus macaques (4, 10), while enteropathogenic *E. coli* strains can—in the laboratory—cause colitis in marmosets (11), rhesus macaques infected with simian immunodeficiency virus (12) and cotton-top tamarins (13).

There are two potential explanations for the co-occurrence of *E. coli* in humans and non-human primates. Some bacterial lineages may have been passed on through vertical transmission within the same host species for long periods, perhaps even arising from ancestral bacteria that colonised the guts of the most recent common ancestors of humans and non-human primate species (14–16). In such a scenario, isolates from non-human primates would be expected to be novel and distinct from the diversity seen in humans. However, there is also clearly potential for horizontal transfer of strains from one host species to another (17).

The exchange of bacteria between humans and human-habituated animals, particularly non-human primates, is of interest in light of the fragmentation of natural habitats globally (18–28). We have seen that wild non-human primates in the Gambia are frequently exposed to humans through tourism, deforestation and urbanisation. In Uganda, PCR-based studies have suggested transmission of *E. coli* between humans, non-human primates and livestock (26–28). Thus, wild non-human primates may constitute a reservoir for the zoonotic spread of *E. coli* strains associated with virulence and antimicrobial resistance to humans. Alternatively, humans might provide a reservoir of strains with the potential for anthroponotic spread to animals—or transmission might occur in both directions (29).

We do not know how many different lineages can co-exist within the same non-human primate host. Such information may help us contextualise the potential risks associated with transmission of bacterial strains between humans and non-human primates. In humans, up to eleven serotypes could be sampled from picking eleven colonies from individual stool samples (30).

To address these issues, we have exploited whole-genome sequencing to explore the colonisation patterns, population structure and phylogenomic diversity of *E. coli* in wild non-human primates from rural and urban Gambia.

## Methods

### Study population and sample collection

In June 2017, wild non-human primates were sampled from six sampling sites in the Gambia: Abuko Nature Reserve (riparian forest), Bijilo Forest Park (coastal fenced woodland), Kartong village (mangrove swamp), Kiang West National park (dry-broad-leaf forest), Makasutu Cultural Forest (ecotourism woodland) and River Gambia National park (riparian forest) (Figure 1). We sampled all four of the diurnal non-human primate species indigenous to the Gambia. Monkeys in Abuko and Bijilo are frequently hand-fed by visiting tourists, despite prohibiting guidelines (31).

**Figure 1.**
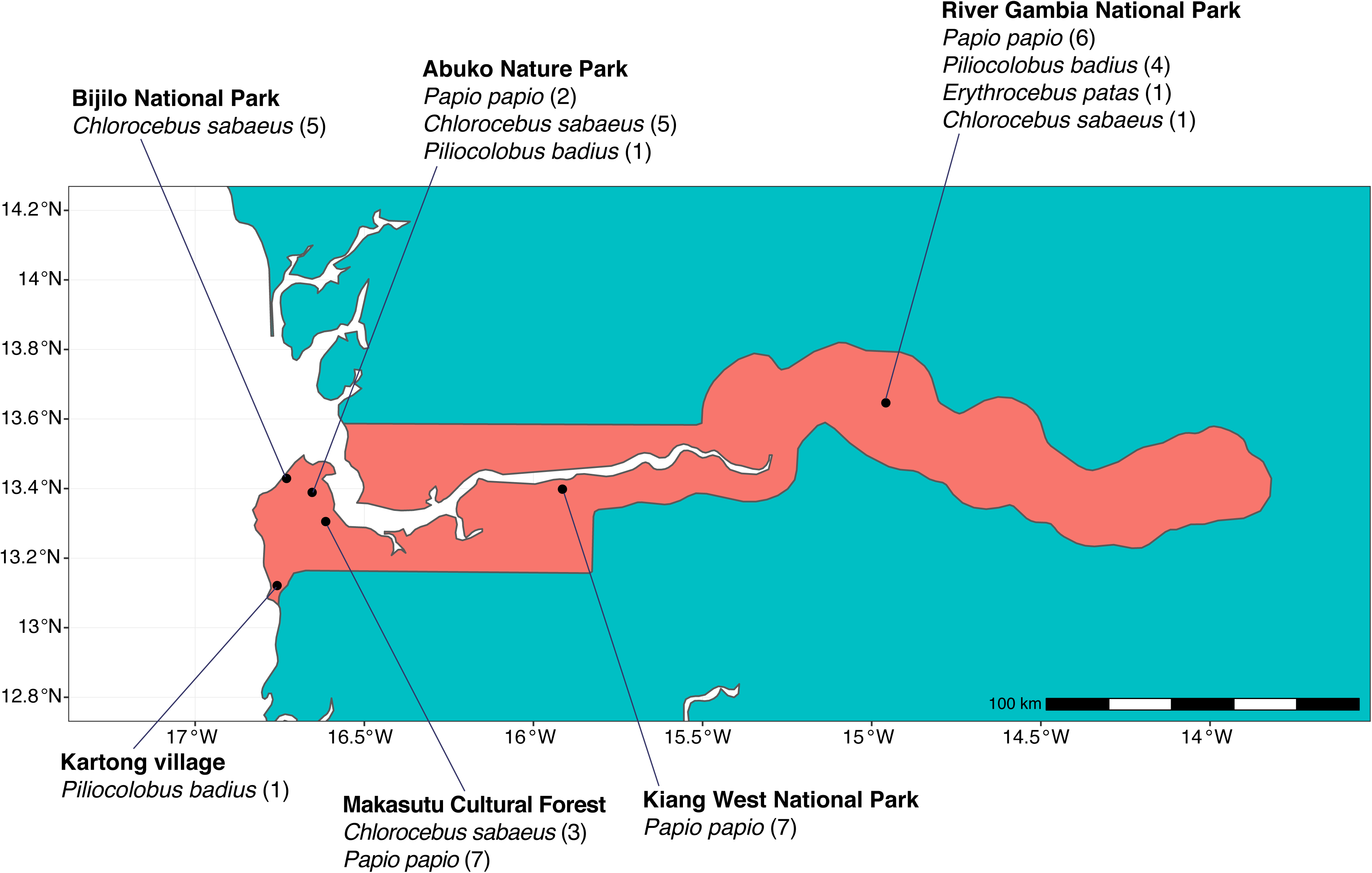
Study sites and distribution of study subjects.

Troops of monkeys were observed and followed. We collected a single freshly passed formed stool specimen from 43 visibly healthy individuals (38 adults, 5 juveniles; 24 females, 11 males, 8 of undetermined sex), drawn from four species: *Erythrocebus patas* (patas monkey), *Papio papio* (Guinea baboon), *Chlorocebus sabaeus* (green monkey) and *Piliocolobus badius* (Western colobus monkey). Stool samples were immediately placed into sterile falcon tubes, taking care to collect portions of stool material that had not touched the ground, then placed on dry ice and stored at 80°C within 6 h. The sample processing flow is summarised in Figure 2.

**Figure 2.**
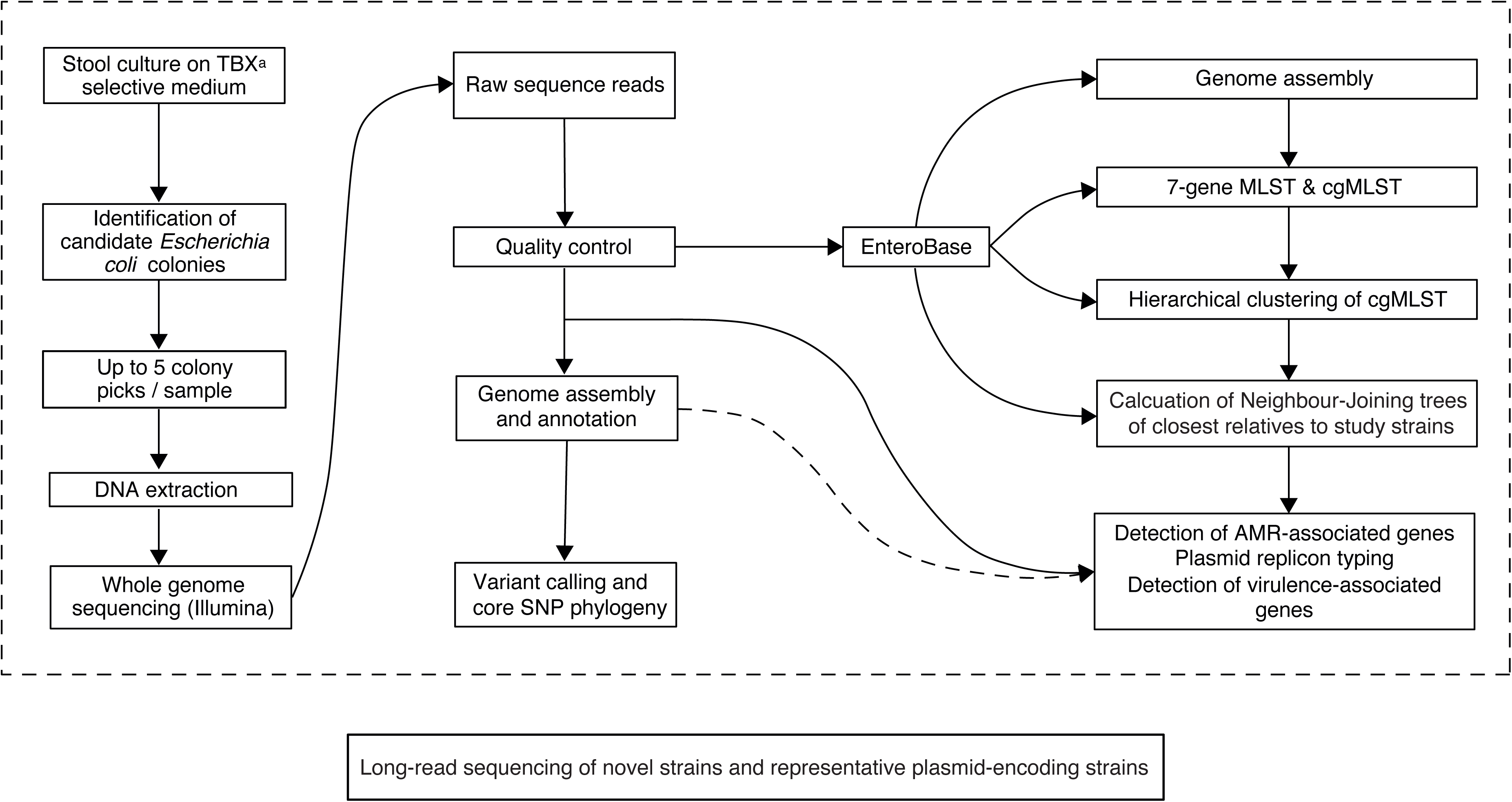
Study sample-processing flow diagram.

### Microbiological processing

For the growth and isolation of *E. coli*, 0.1–0.2 g aliquots were taken from each stool sample into 1.5 ml microcentrifuge tubes under aseptic conditions. To each tube, 1 ml of physiological saline (0.85%) was added, and the saline-stool samples were vortexed for 2 min at 4200 rpm. The homogenised samples were taken through four ten-fold serial dilutions and a 100 µl aliquot from each dilution was spread on a plate of tryptone-bile-X-glucoronide agar using the cross-hatching method. Plates were incubated at 37°C for 18–24 h in air. Colony counts were performed for each serial dilution, counting translucent colonies with blue-green pigmentation and entire margins as *E. coli*. Up to five colonies from each sample were sub-cultured on MacConkey agar at 37°C for 18–24 h and then stored in 20% glycerol broth at −80°C.

### Genomic DNA extraction

A single colony from each subculture was picked into 1 ml Luria-Bertani broth and incubated overnight at 37°C. Broth cultures were spun at 3500rpm for 2 min and lysed using lysozyme, proteinase K, 10% SDS and RNase A in Tris EDTA buffer (pH 8.0). Suspensions were placed on a thermomixer with vigorous shaking at 1600 rpm, first at 37°C for 25 min and subsequently at 65°C for 15 min. DNA was extracted using solid-phase reversible immobilisation magnetic beads (Becter Coulter Inc., Brea, CA, U.S.A.), precipitated with ethanol, eluted in Tris-Cl and evaluated for protein and RNA contamination using A_260_/A_280_ and A_260_/A_230_ ratios on the NanoDrop 2000 Spectrophotometer (Fisher Scientific, Loughborough, UK). DNA concentrations were measured using the Qubit HS DNA assay (Invitrogen, MA, USA). DNA was stored at −20°C.

### Illumina sequencing

Whole-genome sequencing was carried out on the Illumina NextSeq 500 platform (Illumina, San Diego, CA). We used a modified Nextera XT DNA protocol for the library preparation as follows. The genomic DNA was normalised to 0.5 ng µl^-1^ with 10 mM Tris-HCl. Next, 0.9 µl of Tagment DNA buffer (Illumina Catalogue No. 15027866) was mixed with 0.09 µl of Tagment DNA enzyme (Illumina Catalogue No. 15027865) and 2.01 µl of PCR-grade water in a master-mix. Next, 3 µl of the master-mix was added to a chilled 96-well plate. To this, 2 µl of normalised DNA (1 ng total) was added, pipette-mixed and the reaction heated to 55°C for 10 min on a PCR block. To each well, we added 11 µl of KAPA2G Robust PCR master-mix (Sigma Catalogue No. KK5005), comprising 4 µl KAPA2G buffer, 0.4 µl dNTPs, 0.08 µl polymerase and 6.52 µl PCR-grade water, contained in the kit per sample. Next, 2 µl each of P7 and P5 Nextera XT Index Kit v2 index primers (Illumina Catalogue numbers FC-131-2001 to 2004) were added to each well. Finally, the 5 µl of Tagmentation mix was added and mixed. The PCR was run as follows: 72°C for 3 min, 95°C for 1 min, 14 cycles of 95°C for 10 sec, 55°C for 20 sec and 72°C for 3 min. Following the PCR, the libraries were quantified using the Quant-iT dsDNA Assay Kit, high sensitivity kit (Catalogue No. 10164582) and run on a FLUOstar Optima plate reader. After quantification, libraries were pooled in equal quantities. The final pool was double-SPRI size-selected between 0.5 and 0.7x bead volumes using KAPA Pure Beads (Roche Catalogue No. 07983298001). We then quantified the final pool on a Qubit 3.0 instrument (Invitrogen, MA, USA) and ran it on a high sensitivity D1000 ScreenTape (Agilent Catalogue No. 5067-5579) using the Agilent TapeStation 4200 to calculate the final library pool molarity. The pooled library was run at a final concentration of 1.8 pM on an Illumina NextSeq500 instrument using a mid-output flow cell (NSQ® 500 Mid Output KT v2 300 cycles; Illumina Catalogue No. FC-404-2003) following the Illumina recommended denaturation and loading parameters, which included a 1% PhiX spike (PhiX Control v3; Illumina Catalogue FC-110-3001). The data was uploaded to BaseSpace (http://www.basespace.illumina.com) and then converted to FASTQ files.

### Oxford nanopore sequencing

We used the rapid barcoding kit (Oxford Nanopore Catalogue No. SQK-RBK004) to prepare libraries according to the manufacturer’s instructions. We used 400 ng DNA for library preparation and loaded 75 µl of the prepared library on an R9.4 MinION flow cell. The size of the DNA fragments was assessed using the Agilent 2200 TapeStation (Agilent Catalogue No. 5067-5579) before sequencing. The concentration of the final library pool was measured using the Qubit high-sensitivity DNA assay (Invitrogen, MA, USA).

### Genome assembly and phylogenetic analysis

Sequences were analysed on the Cloud Infrastructure for Microbial Bioinformatics (32). Paired-end short-read sequences were concatenated, then quality-checked using FastQC v0.11.7 (33). Reads were assembled using Shovill (https://github.com/tseemann/shovill) and assemblies assessed using QUAST v 5.0.0, de6973bb (34). Draft bacterial genomes were annotated using Prokka v 1.13 (35). Multi-locus sequence types were called from assemblies according to the Achtman scheme using the mlst software (https://github.com/tseemann/mlst) to scan alleles in PubMLST (https://pubmlst.org/) (36). To identify and assign new STs, we used the ST search algorithm in EnteroBase, allowing for one allele mismatch (37). Snippy v4.3.2 (https://github.com/tseemann/snippy) was used for variant calling and core genome alignment, including references genome sequences representing the major phylogroups of *E. coli* and *Escherichia fergusonii* as an outgroup (Supplementary File 1B). We used Gubbins (Genealogies Unbiased By recomBinations In Nucleotide Sequences) to detect and remove recombinant regions of the core genome alignment (38). RAxML v 8.2.4 (39) was used for maximum-likelihood phylogenetic inference from this masked alignment based on a general time-reversible nucleotide substitution model with 1,000 bootstrap replicates. The phylogenetic tree was visualised using Mega v. 7.2 (40) and annotated using Adobe Illustrator v 23.0.3 (Adobe Inc., San Jose, California). Pair-wise single nucleotide polymorphism (SNP) distances between genomes were computed from the core-gene alignment using snp-dists v0.6 (https://github.com/tseemann/snp-dists).

### Population structure and analysis of gene content

Merged short reads were uploaded to EnteroBase (41) where we used the Hierarchical Clustering (HierCC) algorithm to assign our genomes from non-human primates to HC1100 clusters, which in *E. coli* correspond roughly to the clonal complexes seen in seven-allele MLST. Core genome MLST (cgMLST) profiles based on the typing of 2, 512 core loci for *E. coli* facilitates single-linkage hierarchical clustering according to fixed core genome MLST (cgMLST) allelic distances, based on cgMLST allelic differences. Thus, cgST HierCC provides a robust approach to analyse population structures at multiple levels of resolution. The identification of closely-related genomes using HierCC has been shown to be 89% consistent between cgMLST and single-nucleotide polymorphisms (42). Neighbour-joining trees were reconstructed with Ninja—a hierarchical clustering algorithm for inferring phylogenies that is capable of scaling to inputs larger than 100,000 sequences (43).

ARIBA v2.12.1 (44) was used to search short reads against the Virulence Factors Database (45) (VFDB-core) (virulence-associated genes), ResFinder (AMR) (46) and PlasmidFinder (plasmid-associated genes) (47) databases (both ResFinder and PlasmidFinder databases downloaded 29 October 2018). Percentage identity of ≥ 90% and coverage of ≥ 70% of the respective gene length were taken as a positive result. Analyses were performed on assemblies using ABRicate v 0.8.7 (https://github.com/tseemann/abricate). A heat map of detected virulence- and AMR-associated genes was plotted on the phylogenetic tree using ggtree and phangorn in R studio v 3.5.1. We searched EnteroBase for all *E. coli* strains isolated from humans in the Gambia (n=128), downloaded the genomes and screened them for resistance genes using ABRicate v 0.9.8. Assembled genomes for isolates that clustered with our colibactin-encoding ST73, ST127 and ST681 isolates were downloaded and screened for the colibactin operon using ABRicate’s VFDB database (accessed 28 July 2019). Assemblies reported to contain colibactin genes were aligned against the colibactin-encoding *Escherichia coli* IHE3034 reference genome (NCBI Accession: GCA_000025745.1) using minimap2 2.13-r850. BAM files were visualised in Artemis Release 17.0.1 (48) to confirm the presence of the *pks* genomic island which encodes the colibactin operon.

### Hybrid assembly and analysis of plasmids and phages

Base-called FASTQ files were concatenated into a single file and demultiplexed into individual FASTQ files based on barcodes, using the qcat python command-line tool v 1.1.0 (https://github.com/nanoporetech/qcat). Hybrid assemblies of the Illumina and nanopore reads were created with Unicycler (49). The quality and completion of the hybrid assemblies were assessed with QUAST v 5.0.0, de6973bb and CheckM (34, 50). Hybrid assemblies were interrogated using ABRicate PlasmidFinder and annotated using Prokka (35). Plasmid sequences were visualised in Artemis using coordinates from ABRicate. Prophage identification was carried out using the phage search tool, PHASTER (51).

### Antimicrobial susceptibility

We determined the minimum inhibitory concentrations of amikacin, trimethoprim, sulfamethoxazole, ciprofloxacin, cefotaxime and tetracycline for the isolates from non-human primates using agar dilution (52). Two-fold serial dilutions of each antibiotic were performed in molten Mueller-Hinton agar (Oxoid, Basingstoke, UK), from 32mg/L to 0.03 mg l^-1^ (512 mg l^-1^ to 0.03 mg l^-1^ for sulfamethoxazole), using *E. coli* NCTC 10418 as control. MICs were performed in duplicate and interpreted using breakpoint tables from the European Committee on Antimicrobial Susceptibility Testing v. 9.0, 2019 (http://www.eucast.org).

## Results

Twenty-four of 43 samples (56%) showed growth indicative of *E. coli*, yielding a total of 106 colonies. The isolates were designated by the primate species and the site from which they were sampled as follows: *Chlorocebus sabaeus*, ‘Chlos’; *Papio papio*, ‘Pap’; *Piliocolobus badius*, ‘Prob’; Abuko Nature Reserve, ‘AN’; Bijilo Forest Park, ‘BP’; Kartong village, ‘K’; Kiang West National Park, ‘KW’; Makasutu Cultural Forest, ‘M’; and River Gambia National Park, ‘RG’. After genome sequencing, five isolates (PapRG-04, (n=1); PapRG-03 (n=1); ChlosRG-12 (n=1); ChlosAN-13 (n=1); ProbAN-19 (n=1)) were excluded due to low depth of coverage (<20x), leaving 101 genomes for subsequent analysis (Table 1).

**Table 1:** Study isolates.

| Name | Source | Individual sampling number | Colony-pick | Sampling site | ST |
| --- | --- | --- | --- | --- | --- |
| PapRG-03-1 | *Papio papio* | 3 | 1 | River Gambia national park | 336 |
| PapRG-03-2 | *Papio papio* | 3 | 2 | River Gambia national park | 336 |
| PapRG-03-3 | *Papio papio* | 3 | 3 | River Gambia national park | 336 |
| PapRG-03-4 | *Papio papio* | 3 | 4 | River Gambia national park | 336 |
| PapRG-03-5 | *Papio papio* | 3 | 5 | River Gambia national park | 336 |
| PapRG-04-1 | *Papio papio* | 4 | 1 | River Gambia national park | 1665 |
| PapRG-04-2 | *Papio papio* | 4 | 2 | River Gambia national park | 1204 |
| PapRG-04-4 | *Papio papio* | 4 | 3 | Makasutu cultural forest | 8826 |
| PapRG-04-5 | *Papio papio* | 4 | 4 | Makasutu cultural forest | 1204 |
| PapRG-05-2 | *Papio papio* | 5 | 1 | Makasutu cultural forest | 1431 |
| PapRG-05-3 | *Papio papio* | 5 | 2 | Makasutu cultural forest | 99 |
| PapRG-05-4 | *Papio papio* | 5 | 3 | Makasutu cultural forest | 6316 |
| PapRG-05-5 | *Papio papio* | 5 | 4 | Makasutu cultural forest | 1431 |
| PapRG-06-1 | *Papio papio* | 6 | 1 | Makasutu cultural forest | 4080 |
| PapRG-06-2 | *Papio papio* | 6 | 2 | Makasutu cultural forest | 2521 |
| PapRG-06-3 | *Papio papio* | 6 | 3 | Makasutu cultural forest | 8827 |
| PapRG-06-4 | *Papio papio* | 6 | 4 | Makasutu cultural forest | 1204 |
| PapRG-06-5 | *Papio papio* | 6 | 5 | River Gambia national park | 8525 |
| ProbRG-07-1 | *Piliocolobus badius* | 7 | 1 | River Gambia national park | 73 |
| ProbRG-07-2 | *Piliocolobus badius* | 7 | 2 | River Gambia national park | 73 |
| ProbRG-07-3 | *Piliocolobus badius* | 7 | 3 | River Gambia national park | 73 |
| ProbRG-07-4 | *Piliocolobus badius* | 7 | 4 | River Gambia national park | 73 |
| ProbRG-07-5 | *Piliocolobus badius* | 7 | 5 | River Gambia national park | 73 |
| ChlosRG-12-1 | *Chlorocebus sabaesus* | 12 | 1 | River Gambia national park | 8824 |
| ChlosRG-12-2 | *Chlorocebus sabaesus* | 12 | 2 | River Gambia national park | 196 |
| ChlosRG-12-3 | *Chlorocebus sabaesus* | 12 | 3 | River Gambia national park | 196 |
| ChlosRG-12-5 | *Chlorocebus sabaesus* | 12 | 4 | River Gambia national park | 40 |
| ChlosAN-13-1 | *Chlorocebus sabaesus* | 13 | 1 | Abuko Nature Reserve | 8526 |
| ChlosAN-13-2 | *Chlorocebus sabaesus* | 13 | 2 | Abuko Nature Reserve | 8550 |
| ChlosAN-13-4 | *Chlorocebus sabaesus* | 13 | 3 | Abuko Nature Reserve | 1973 |
| ChlosAN-13-5 | *Chlorocebus sabaesus* | 13 | 4 | Abuko Nature Reserve | 1973 |
| PapAN-14-1 | *Papio papio* | 14 | 1 | Abuko Nature Reserve | 2076 |
| PapAN-14-2 | *Papio papio* | 14 | 2 | Abuko Nature Reserve | 939 |
| PapAN-14-3 | *Papio papio* | 14 | 3 | Abuko Nature Reserve | 226 |
| PapAN-14-4 | *Papio papio* | 14 | 4 | Abuko Nature Reserve | 226 |
| PapAN-14-5 | *Papio papio* | 14 | 5 | Abuko Nature Reserve | 226 |
| PapAN-15-1 | *Papio papio* | 15 | 1 | Abuko Nature Reserve | 226 |
| PapAN-15-2 | *Papio papio* | 15 | 2 | Abuko Nature Reserve | 5073 |
| PapAN-15-3 | *Papio papio* | 15 | 3 | Abuko Nature Reserve | 226 |
| PapAN-15-4 | *Papio papio* | 15 | 4 | Abuko Nature Reserve | 126 |
| PapAN-15-5 | *Papio papio* | 15 | 5 | Abuko Nature Reserve | 8823 |
| ChlosAN-17-1 | *Chlorocebus sabaesus* | 17 | 1 | Abuko Nature Reserve | 681 |
| ChlosAN-17-2 | *Chlorocebus sabaesus* | 17 | 2 | Abuko Nature Reserve | 362 |
| ChlosAN-17-3 | *Chlorocebus sabaesus* | 17 | 3 | Abuko Nature Reserve | 681 |
| ChlosAN-17-4 | *Chlorocebus sabaesus* | 17 | 4 | Abuko Nature Reserve | 681 |
| ChlosAN-18-1 | *Chlorocebus sabaesus* | 18 | 1 | Abuko Nature Reserve | 681 |
| ChlosAN-18-2 | *Chlorocebus sabaesus* | 18 | 2 | Abuko Nature Reserve | 681 |
| ChlosAN-18-3 | *Chlorocebus sabaesus* | 18 | 3 | Abuko Nature Reserve | 681 |
| ChlosAN-18-4 | *Chlorocebus sabaesus* | 18 | 4 | Abuko Nature Reserve | 681 |
| ChlosAN-18-5 | *Chlorocebus sabaesus* | 18 | 5 | Abuko Nature Reserve | 349 |
| ProbAN-19-2 | *Ptilocolobus badius* | 19 | 1 | Abuko Nature Reserve | 8825 |
| ChlosBP-21-1 | *Chlorocebus sabaesus* | 21 | 1 | Bijilo forest park | 677 |
| ChlosBP-21-2 | *Chlorocebus sabaesus* | 21 | 2 | Bijilo forest park | 677 |
| ChlosBP-21-3 | *Chlorocebus sabaesus* | 21 | 3 | Bijilo forest park | 677 |
| ChlosBP-21-4 | *Chlorocebus sabaesus* | 21 | 4 | Bijilo forest park | 677 |
| ChlosBP-21-5 | *Chlorocebus sabaesus* | 21 | 5 | Bijilo forest park | 677 |
| ChlosBP-23-1 | *Chlorocebus sabaesus* | 23 | 2 | Bijilo forest park | 8527 |
| ChlosBP-23-2 | *Chlorocebus sabaesus* | 23 | 3 | Bijilo forest park | 8527 |
| ChlosBP-23-3 | *Chlorocebus sabaesus* | 23 | 4 | Bijilo forest park | 3306 |
| ChlosBP-24-1 | *Chlorocebus sabaesus* | 24 | 1 | Bijilo forest park | 73 |
| ChlosBP-24-2 | *Chlorocebus sabaesus* | 24 | 2 | Bijilo forest park | 73 |
| ChlosBP-24-3 | *Chlorocebus sabaesus* | 24 | 3 | Bijilo forest park | 73 |
| ChlosBP-24-4 | *Chlorocebus sabaesus* | 24 | 4 | Bijilo forest park | 73 |
| ChlosBP-24-5 | *Chlorocebus sabaesus* | 24 | 5 | Bijilo forest park | 73 |
| ChlosBP-25-1 | *Chlorocebus sabaesus* | 25 | 1 | Bijilo forest park | 73 |
| ChlosBP-25-2 | *Chlorocebus sabaesus* | 25 | 2 | Bijilo forest park | 73 |
| ChlosBP-25-3 | *Chlorocebus sabaesus* | 25 | 3 | Bijilo forest park | 73 |
| ChlosBP-25-4 | *Chlorocebus sabaesus* | 25 | 4 | Bijilo forest park | 73 |
| ChlosBP-25-5 | *Chlorocebus sabaesus* | 25 | 5 | Bijilo forest park | 73 |
| ChlosM-29-1 | *Chlorocebus sabaesus* | 29 | 1 | Makasutu cultural forest | 1873 |
| ChlosM-29-2 | *Chlorocebus sabaesus* | 29 | 2 | Makasutu cultural forest | 1873 |
| PapM-31-1 | *Papio papio* | 31 | 1 | Makasutu cultural forest | 2800 |
| PapM-31-2 | *Papio papio* | 31 | 2 | Makasutu cultural forest | 135 |
| PapM-31-3 | *Papio papio* | 31 | 3 | Makasutu cultural forest | 5780 |
| PapM-31-4 | *Papio papio* | 31 | 4 | Makasutu cultural forest | 1727 |
| PapM-31-5 | *Papio papio* | 31 | 5 | Makasutu cultural forest | 5780 |
| PapM-32-1 | *Papio papio* | 32 | 2 | Makasutu cultural forest | 8532 |
| PapM-32-2 | *Papio papio* | 32 | 3 | Makasutu cultural forest | 212 |
| PapM-32-3 | *Papio papio* | 32 | 4 | Makasutu cultural forest | 212 |
| PapM-32-4 | *Papio papio* | 32 | 5 | Makasutu cultural forest | 212 |
| PapM-32-5 | *Papio papio* | 32 | 6 | Makasutu cultural forest | 212 |
| PapM-33-1 | *Papio papio* | 33 | 1 | Makasutu cultural forest | 8533 |
| PapM-33-2 | *Papio papio* | 33 | 2 | Makasutu cultural forest | 8533 |
| PapM-33-3 | *Papio papio* | 33 | 3 | Makasutu cultural forest | 8533 |
| PapM-33-4 | *Papio papio* | 33 | 4 | Makasutu cultural forest | 38 |
| PapM-33-5 | *Papio papio* | 33 | 5 | Makasutu cultural forest | 8533 |
| PapM-34-1 | *Papio papio* | 34 | 1 | Makasutu cultural forest | 676 |
| PapM-34-2 | *Papio papio* | 34 | 2 | Makasutu cultural forest | 676 |
| PapM-34-3 | *Papio papio* | 34 | 3 | Makasutu cultural forest | 676 |
| PapM-34-4 | *Papio papio* | 34 | 4 | Makasutu cultural forest | 676 |
| PapM-36-1 | *Papio papio* | 36 | 1 | Makasutu cultural forest | 8535 |
| PapM-36-2 | *Papio papio* | 36 | 2 | Makasutu cultural forest | 8535 |
| PapKW-44-1 | *Papio papio* | 44 | 1 | Kiang West national park | 442 |
| PapKW-44-2 | *Papio papio* | 44 | 2 | Kiang West national park | 442 |
| PapKW-44-3 | *Papio papio* | 44 | 3 | Kiang West national park | 442 |
| PapKW-44-4 | *Papio papio* | 44 | 4 | Kiang West national park | 442 |
| ProbK-45-1 | *Ptilocolobus badius* | 45 | 1 | Kartong village | 127 |
| ProbK-45-2 | *Ptilocolobus badius* | 45 | 2 | Kartong village | 127 |
| ProbK-45-3 | *Ptilocolobus badius* | 45 | 3 | Kartong village | 127 |
| ProbK-45-4 | *Ptilocolobus badius* | 45 | 4 | Kartong village | 127 |
| ProbK-45-5 | *Ptilocolobus badius* | 45 | 5 | Kartong village | 127 |

We recovered 43 seven-allele sequence types (ten of them novel), spanning five of the eight known phylogroups of *E. coli* and comprising 38 core-genome MLST complexes (Figure 3). The majority of strains belonged to phylogroup B2 (42/101, 42%), which encompasses strains that cause extraintestinal infections in humans (ExPEC strains) (6–8). Strains from phylogroup B2 carried colonisation and fitness factors associated with extraintestinal disease in humans (Figure 3). A subset of the B2 strains (13/42, 31%), belonging to STs 73, 681 and 127, carried the *pks* genomic island, which encodes the DNA alkylating genotoxin, colibactin. Colibactin-encoding *E. coli* frequently cause colorectal cancer, urosepsis, bacteraemia and prostatitis, and are highly associated with other virulence factors such as siderophores and toxins (53–56).

**Figure 3.**
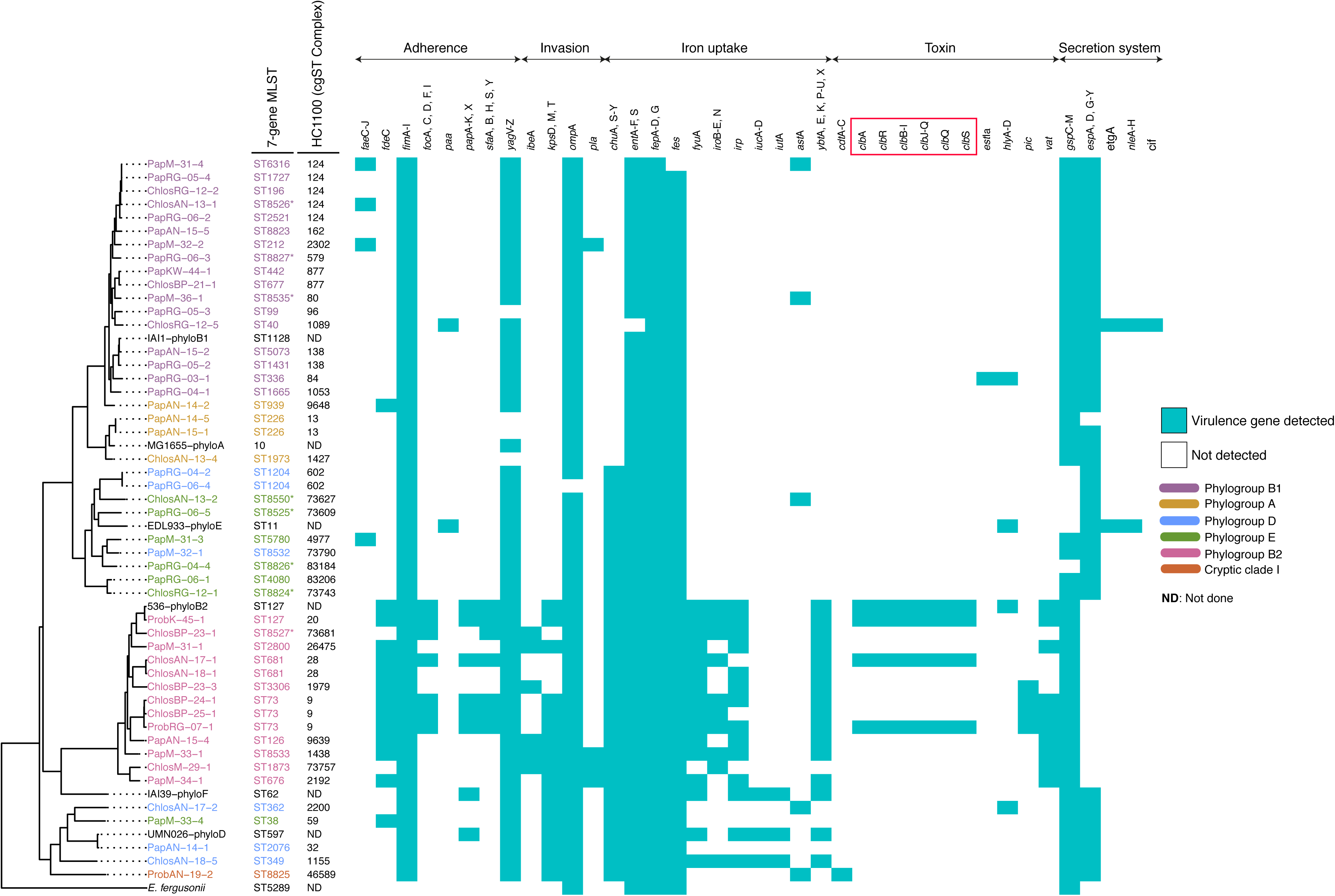
A plot showing the maximum likelihood phylogeny of the study isolates overlaid with the prevalence of potential virulence genes among the study isolates. The tree was reconstructed based on non-repetitive core SNPs calculated against the *E. coli* K-12 reference strain (NCBI accession: NC_000913.3), using RAxML with 1000 bootstrap replicates. *E. coli* MG1655 was used as the reference and *E. fergusonii* as the outroot species. Recombinant regions were removed using Gubbins (Reference 38). The tip labels indicate the sample IDs, with the respective in silico Achtman sequence types (STs) and HC1100 (cgST complexes) are indicated next to the tip labels. Both the sample IDs and the STs (Achtman) are colour-coded to indicate the various phylogroups as indicated. Novel STs (Achtman) are indicated by an asterisk (*). *Escherichia fergusonii* and the *E. coli* reference genomes representing the major *E. coli* phylogroups are in black. Primate species are indicated in the strain names as follows: *Chlorocebus sabaeus*, ‘Chlos’; *Papio papio*, ‘Pap’; *Piliocolobus badius*, ‘Prob’. The sampling sites are indicated as follows: BP, Bijilo forest park; KW, Kiang-West National park; RG, River Gambia National Park; M, Makasutu Cultural forest; AN, Abuko Nature reserve; K, Kartong village. Co-colonising seven-allele (Achtman) sequence types (STs) in single individuals are shown by the prefix of the strain names depicting the colony as 1, 2 up to 5. We do not show multiple colonies of the same Achtman ST recovered from a single individual. In such cases, only one representative is shown. Virulence genes are grouped according to their function, with genes encoding the colibactin genotoxin highlighted with a red box. The full names of virulence factors are provided in Supplementary file 5.

Thirteen individuals were colonised by two or more STs and nine by two or more phylogroups (Supplementary File 1A). Five colony picks from a single Guinea baboon (PapRG-06) yielded five distinct STs, two of which are novel. Two green monkeys sampled from Bijilo (ChlosBP-24 and ChlosBP-25) shared an identical ST73 genotype, while two Guinea baboons from Abuko shared an ST226 strain—documenting transmission between monkeys of the same species. Among the monkey isolates, we found several STs associated with extraintestinal infections and/or AMR in humans: ST73, ST681, ST127, ST226, ST336, ST349 (57–62).

In seventeen monkeys, we observed a cloud of closely related genotypes (sepearated by 0-5 SNPs, Table 2A) from each strain, suggesting evolution within the host after acqusition of the strain. However, in two individuals, pair-wise SNP distances between genotypes from the same ST were susbtantial enough (25 SNPs and 77 SNPs) to suggest multiple acquisitions of each strain (Table 2B).

**Table 2A:** Within-host single nucleotide polymorphism diversity between multiple genomes of the same ST recovered from the same monkey.

| Sample ID | STs (colonies per ST) | Pair-wise SNP distances between multiple colonies of the same ST | Comment(s) |
| --- | --- | --- | --- |
| PapRG-03 | 336 (n=5) | 0-2 |  |
| PapRG-04 | 1204 (n=2) | 4 |  |
| PapRG-05 | 1431 (n=2) | 0 |  |
| ProbRG-07 | 73 (n=5) | 0-1 |  |
| ChlosRG-12 | 196 (n=2) | 25 |  |
| PapAN-14 | 226 (n=3) | 1 |  |
| PapAN-15 | 226 (n=2) | 1 |  |
| ChlosAN-17 | 681 (n=3) | 0-3 |  |
| ChlosAN-18 | 681 (n=4) | 0 |  |
| ChlosBP-21 | 677 (n=4) | 5 |  |
| ChlosBP-23 | 8527 (n=2) | 0 |  |
| ChlosBP-24 | 73 (n=5) | 0-5 |  |
| ChlosBP-25 | 73 (n=5) | 0-79 | Please see Table 2B |
| PapM-32 | 212 (n=4) | 0 |  |
| PapM-33 | 8533 (n=4) | 0-4 |  |
| PapM-34 | 676 (n=4) | 0-1 |  |
| PapM-36 | 8535 (n=2) | 0-1 |  |
| PapKW-44 | 442 (n=4) | 1-2 |  |
| ProbK-45 | 127 (n=5) | 0-4 |  |

**Table 2B:**
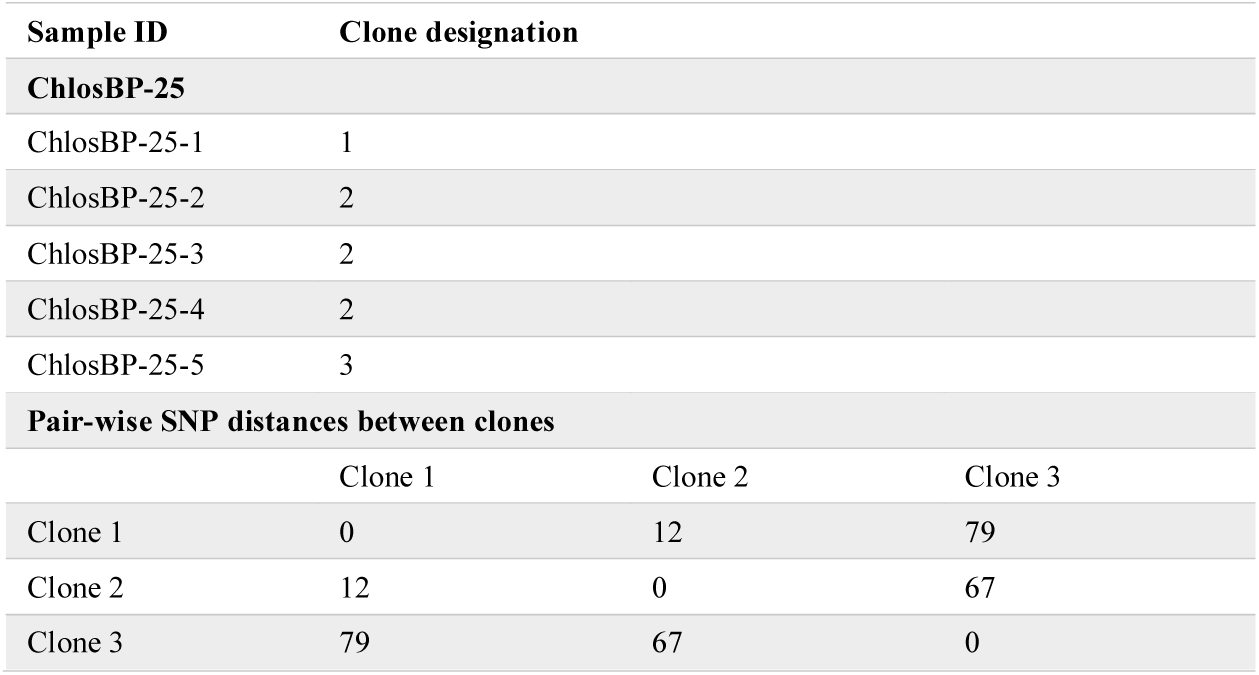
Within-host diversity in green monkey 25 (ChlosBP-25)

We identified the closest neighbours to all the recovered strains from our study (Table 3). Our results suggest, in some cases, recent interactions between humans or livestock and non-human primates. However, we also found a diversity of strains specific to the non-human primate niche. Hierarchical clustering analysis revealed that simian isolates from ST442 and ST349 (Achtman)— sequence types that are associated with virulence and AMR in humans (49, 55)—were closely related to human clinical isolates, with differences of 50 alleles and seven alleles in the core-genome MLST scheme respectively (Supplementary Figures 1-2). Similarly, we found evidence of recent interaction between simian ST939 isolates and strains from livestock (Supplementary Figure 3). Conversely, simian ST73, ST127 and ST681 isolates were genetically distinct from human isolates from these sequence types (Supplementary Figures 4-6). The multi-drug resistant isolate PapAN-14-1 from ST349 was, however, closely related to an environmental isolate recovered from water (Supplementary Figure 7).

**Table 3:** Genomic relationship between study isolates and publicly available *E. coli* genomes.

| 7-allele ST | HC100 subgroups | Non-human primate host | Closest neighbours' source | Neighbours' country of isolation | Allelic distance |
| --- | --- | --- | --- | --- | --- |
| 349 | - | *Chlorocebus sabaeus* 18 | Human (bloodstream infection) | Canada | 7 |
| 2076 | - | *Papio papio* 14 | Environment (water) | Unknown | 25 |
| 939 | - | *Papio papio* 14 | Livestock | US | 40 |
| 442 | - | *Papio papio* 44 | Human | China | 50 |
| 2800 | - | *Papio papio* 31 | Unknown | Vietnam | 59 |
| 1973 | - | *Chlorocebus sabaeus* 13 | Unknown | Unknown | 64 |
| 8533 | - | *Papio papio* 33 | Environment (water) | Unknown | 69 |
| 6316 | - | *Papio papio* 05 | Human | Kenya | 97 |
| 1727 | - | *Papio papio* 34 | Human | Kenya | 98 |
| 676 | - | *Papio papio* 34 | Human (bloodstream infection) | UK | 98 |
| 8823 | - | *Papio papio* 15 | Rodent (guinea pig) | Kenya | 101 |
| 1431 | - | *Papio papio* 05 | Human | US | 109 |
| 5073 | - | *Papio papio* 15 | Human | US | 112 |
| 226 | 73641 | *Papio papio* 14 | Human | Tanzania | 112 |
| 8827 | - | *Papio papio* 06 | Human | Unknown | 122 |
| 1204 | 83197 | *Papio papio* 04 | Livestock | Japan | 127 |
| 1204 | 83197 | *Papio papio* 04 | Livestock | Japan | 130 |
| 677 | - | *Chlorocebus sabaeus* 21 | Human | US | 132 |
| 40 | - | *Chlorocebus sabaeus* 12 | Human | UK | 137 |
| 1204 | 83164 | *Papio papio* 06 | Livestock | Japan | 173 |
| 99 | - | *Papio papio* 05 | Human | UK | 180 |
| 362 | - | *Chlorocebus sabaeus* 17 | Food | Kenya | 180 |
| 8825 | - | *Piliocolobus badius* 19 | Human | France | 189 |
| 336 | - | *Papio papio* 03 | Poultry | Kenya | 189 |
| 73 | - | *Chlorocebus sabaeus* 24 | Human | Sweden | 189 |
| 196 | - | *Chlorocebus sabaeus* 12 | Human | Sweden | 197 |
| 2521 | - | *Papio papio* 06 | Livestock | US | 201 |
| 127 | - | *Piliocolobus badius* 45 | Companion animal | US | 229 |
| 681 | - | *ChlosAN* 17 | Human | Norway | 251 |
| 38 | - | *Papio papio* 33 | human | UK | 265 |
| 135 | - | *Papio papio* 31 | Poultry | US | 281 |
| 8824 | - | *Chlorocebus sabaeus* 12 | Environmental* | US | 296 |
| 226 | 100039 | *Papio papio* 14 | Human | Sri Lanka | 318 |
| 8527 | - | *Chlorocebus sabaeus* 23 | Human | Kenya | 323 |
| 8535 | - | *Papio papio* 36 | Environmental (soil) | US | 368 |
| 1665 | - | *Papio papio* 04 | Livestock | UK | 371 |
| 4080 | - | *Papio papio* 06 | Human | Denmark | 507 |
| 8526 | - | *Chlorocebus sabaeus* 13 | Livestock | US | 708 |
| 8532 | - | *Papio papio* 32 | Non-human primate | Gambia (PapM-31-3) | 1102 |
| 8826 | - | *Papio papio* 04 | Livestock | Mozambique | 1255 |
| 8525 | - | *Papio papio* 06 | Livestock/companion animal | Switzerland | 1659 |
| 1873 | - | *Chlorocebus sabaeus* 29 | Environment | US | 1685 |
| 8550 | - | *Chlorocebus sabaeus* 13 | Unknown | Unknown | 2006 |

\*Source details unknown.
Isolates from humans were recovered from stools, except where indicated otherwise.

Five isolates were >1000 alleles away in the core-genome MLST scheme from anything in EnteroBase (Supplementary Figures 8 & 9). Four of these were assigned to novel sequence types in the seven-allele scheme (Achtman) (ST8550, ST8525, ST8532, ST8826), while one belonged to ST1873, which has only two other representatives in EnteroBase: one from a species of wild bird from Australia (*Sericornis frontalis*); the other from water. Besides, ST8550, ST8525, ST8532, ST8826 belonged to novel HierCC 1100 groups (cgST complexes), indicating that they were unrelated to any other publicly available *E. coli* genomes.

We observed few antimicrobial resistance genes in our study population, compared to what prevails in isolates from humans in the Gambia (Figure 4). Phenotypic resistance to single agents was confirmed in ten isolates: to trimethoprim in a single isolate, to sulfamethoxazole in four unrelated isolates and to tetracycline in four closely related isolates from a single animal. A single ST2076 (Achtman) isolate (PapAN-14-1) belonging to the ST349 lineage was resistant to trimethoprim, sulfamethoxazole and tetracycline. The associated resistance genes were harboured on an IncFIB plasmid.

**Figure 4:**
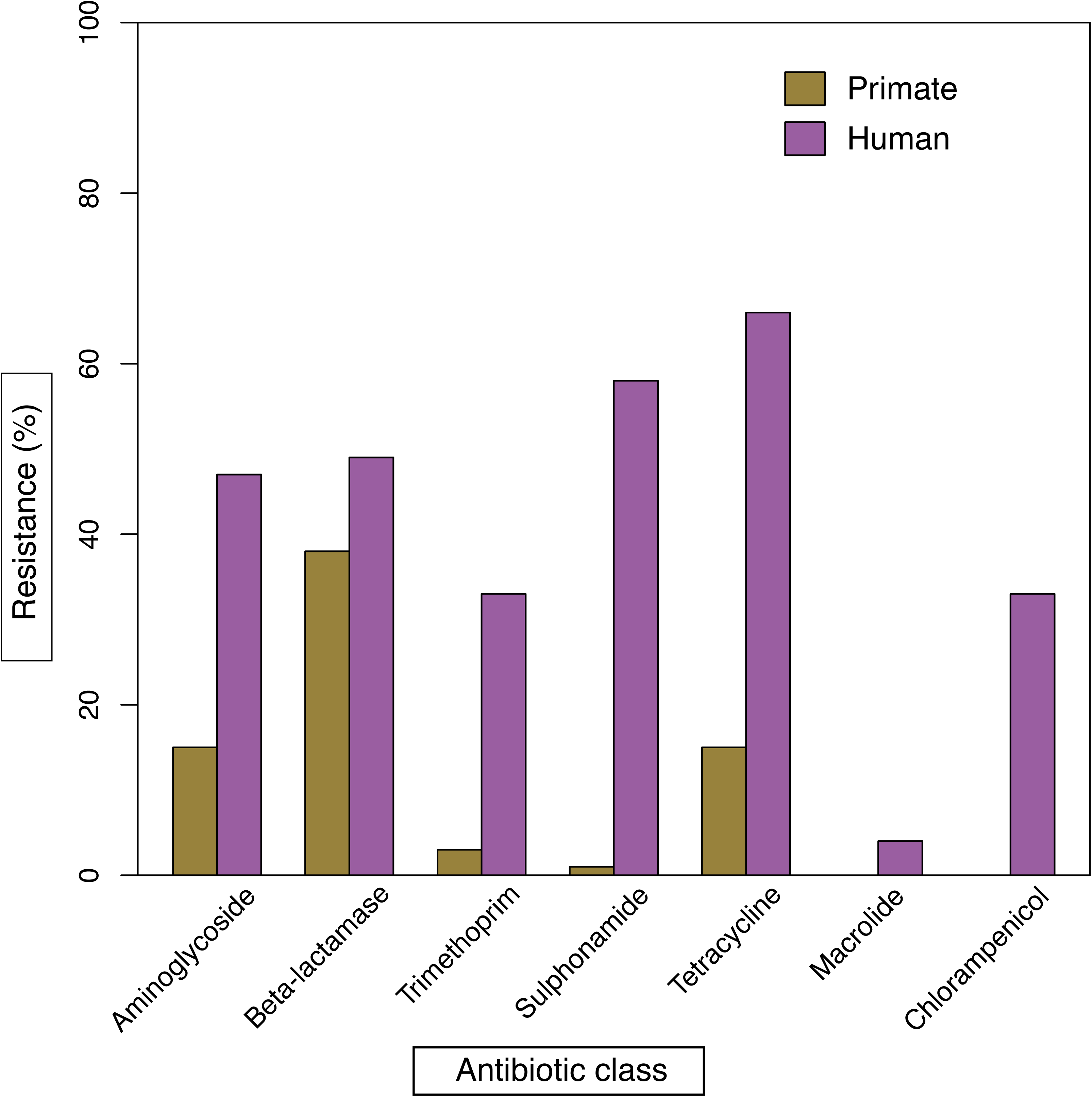
A bar graph comparing the prevalence of antimicrobial resistance genotypes in *E. coli* isolated from humans in the Gambia (n=128) as found in EnteroBase (Reference 41) to that found among the study isolates (n=101). The antimicrobial resistance genes detected were as follows: Aminoglycoside: *aph*(6)-Id, ant *aac*(3)-IIa, *ant*(3’’)-Ia, *aph*(3’’)-Ib, *aad*A1, *aad*A2; Beta-lactamase: *bla*OXA-1, *bla*TEM-1B, *bla*TEM-1B, *bla*TEM-1C, *bla*SHV-1; Trimethoprim: *dfr*A; Sulphonamide: *sul*1, *sul*2; Tetracycline: *tet*(A), *tet*(B), *tet*(34*), tet*(D); Macrolide, *mph*(A); Chloramphenicol*, cat*A1. Screening of resistance genes was carried out using ARIBA ResFinder (Reference 44) and confirmed by ABRicate (https://github.com/tseemann/abricate). A percentage identity of ≥ 90% and coverage of ≥ 70% of the respective gene length were taken as a positive result.

Eighty percent (81/101) of the study isolates harboured one or more plasmids. We detected the following plasmid replicon types: IncF (various subtypes), IncB/K/O/Z, I1, IncX4, IncY, Col plasmids (various subtypes) and plasmids related to p0111 (rep B) (Supplementary File 2A). Long-read sequencing of six representative samples showed that the IncFIB plasmids encoded acquired antibiotic resistance, fimbrial adhesins and colicins (Supplementary File 2B). Also, the IncFIC/FII, ColRNAI, Col156 and IncB/O/K/Z plasmids encoded fimbrial proteins and colicins. Besides, the IncX and Inc-I-Aplha encoded bundle forming pili *bfp*B and the heat-stable enterotoxin protein *StbB* respectively.

We generated complete genome sequences of five novel sequence types of *E. coli* (ST8525, ST8527, ST8532, ST8826, ST8827) within the seven-allele scheme (Achtman) (Supplementary File 3A) (63). Although none of these new genomes encoded AMR genes, ono of them (PapRG-04-4) contained an IncFIB plasmid encoding fimbrial proteins, and a cryptic ColRNA plasmid. PHASTER identified thirteen intact prophages and four incomplete phage remnants (Supplementary File 3B). Two pairs of genomes from Guinea baboons from different parks shared common prophages: one pair carrying PHAGE_Entero_933W, the other PHAGE-Entero_lambda.

## Discussion

We have described the population structure of *E. coli* in diurnal non-human primates living in rural and urban habitats from the Gambia. Although our sample size was relatively small, we have recovered isolates that span the diversity previously described in humans and have also identified ten new sequence types (five of them now with complete genome sequences). This finding is significant, considering the vast number of *E. coli* genomes that have been sequenced to date (9, 597 with MLST via sanger sequencing, and 127, 482 via WGS) (64).

Increasing contact between animal species facilitates the potential exchange of pathogens. Accumulating data shows that ExPEC strains are frequently isolated from diseased companion animals and livestock—highlighting the potential for zoonotic as well as anthroponotic transmission (65–70). In a previous study, green monkeys from Bijilo Park were found to carry lineages of *Staphylococcus aureus* thought to be acquired from humans (31). Our analyses suggest similar exchange of *E. coli* strains between humans and wild non-human primates. However, non-human primates also harbour *E. coli* genotypes that are clinically important in humans, such as ST73, ST127 and ST681, yet are distinct from those circulating in humans—probably reflecting lineages that have existed in this niche for long periods.

We found that several monkeys were colonised with multiple STs, often encompassing two or more phylotypes. Although colonisation with multiple serotypes of *E. coli* is common in humans (30, 71) we were surprised to identify as many as five STs in a single baboon. Sampling multiple colonies from single individuals also revealed within-host diversity arising from microevolution. However, we also found evidence of acquisition in the same animal of multiple lineages of the same sequence type, although it is unclear whether this reflects a single transmission event involving more than one strain or serial transfers.

Antimicrobial resistance in wildlife is known to spread on plasmids through horizontal gene transfer (72). Given the challenge of resolving large plasmids using short-read sequences (73), we exploited long-read sequencing to document the contribution of plasmids to the genomic diversity that we observed in our study population. Consistent with previous reports (74), we found IncF plasmids which encoded antimicrobial resistance genes. Virulence-encoding plasmids, particularly colicin-encoding and the F incompatibility group ones, have long been associated with several pathotypes of *E. coli* (75). Consistent with this, we found plasmids that contributed to the dissemination of virulence factors such as the heat-stable enterotoxin protein *StbB*, colicins and fimbrial proteins.

This study could have been enhanced by sampling human populations living near those of our non-human primates; however, we compensated for this limitation by leveraging the wealth of genomes in publicly available databases. Besides, we did not sample nocturnal monkeys due to logistic challenges; however, these have more limited contact with humans than the diurnal species. Despite these limitations, however, this study provides insight into the diversity and colonisation patterns of *E. coli* among non-human primates in the Gambia, highlighting the impact of human continued encroachment on natural habitats and revealing important phylogenomic relationships between strains from humans and non-human primates.

## Data bibliography

1. Foster-Nyarko, E. et al, NCBI BioProject PRJNA604701 (2020).
2. Forde, B. M., Ben Zakour, N. L., Stanton-Cook, M., Phan, M. D., Totsika, M. et al., 17 representative *E. coli* reference isolates (2014). NCBI accession numbers are provided in Table 1B.
3. Nougayrede J.P, Homburg S, Taieb F., Boury M., Brzuszkiewicz E., et al., *Escherichia coli* induces DNA double-strand breaks in eukaryotic cells (2006). NCBI accession: GCA_000025745.1.

## Acknowledgements

We want to thank Dr Andrew Page and Dr Thanh Le-Viet for their thoughtful advice on the long-read analysis. We also thank Dr Mark Webber for proofreading the manuscript and giving constructive feedback.

## Author contributions

Conceptualization, MA, MP; data curation, MP, NFA; formal analysis, EFN, analytical support, GT; funding, MP and MA; sample collection, JDC; laboratory experiments, EFN, DB; supervision, AR, NFA, GK, JO, MP, MA; manuscript preparation – original draft, EFN; review and editing, NT, AR, JO, NFA, MP; review of final manuscript, all authors.

## Ethical statement

No human nor animal experimentation is reported.

## Figure legends

**Supplementary Figure 1.** A Ninja neighbour-joining tree showing the phylogenetic relationship between Achtman ST442 strains from this study and all other publicly available genomes that fell within the same HC1100 cluster (cgST complex). The locations of the isolates are displayed, with the genome count displayed in parenthesis. Branch lengths display the allelic distances separating genomes. Gambian strains are highlighted in red. The sub-tree (B) shows the closest relatives to the study strains, with the allelic distance separating them displayed with the arrow. Dotted lines represent long branches which have been shortened.

**Supplementary Figure 2.** A Ninja neighbour-joining tree showing the phylogenetic relationship between the ST349 (Achtman) strain from this study and all other publicly available genomes within the same HC1100 cluster (cgST complex). The legend shows the locations of the isolates, with genome counts displayed in parenthesis. Gambian strains are highlighted in red. The study ST349 strain is separated from a clinical ST349 strain by only seven alleles (<7 SNPs), as depicted in the subtree (B). Long branches are shortened (indicated by dashes).

**Supplementary Figure 3.** A phylogenetic neighbour-joining tree reconstructed with the study ST939 (Achtman) strain and all publicly available genomes that fell within the same HC1100 cluster (cgST complex). The legend shows the locations of the isolates, with red highlights around the nodes indicating the Gambian strains. The allelic distance between the study strain and its nearest relative, a bovine ST939 strain, has been given, depicted by the arrow. Dotted lines indicate shortened long branches.

**Supplementary Figure 4.** A Ninja neighbour-joining tree reconstructed with Achtman ST73 colibactin+ strains from this study and all other publicly available ST73 (Achtman) strains that fell within the same HC1100 cluster (cgST complex) in EnteroBase (Reference 41). The sources of the isolates are displayed, with Gambian strains highlighted in red. The Gambian non-human primate strains are on separate long branches, although nested within clades populated by human strains from other countries, suggestive of probably an ancient transmission between the two hosts. The branch lengths for the Gambian strains are displayed. Dotted lines represent long branches which have been shortened.

**Supplementary Figure 5.** A Ninja neighbour-joining tree showing the phylogenetic relationship between ST127 strains from this study and other publicly available strains that occur within the same HC1100 cluster (cgST complex). The sources of the isolates are displayed in the legends, with Gambian strains highlighted in red. Branch lengths display the allelic distances separating genomes. The sub-tree (B) shows the closest relatives to the study strains, with the allelic distances separating them displayed with the arrow. Dotted lines represent long branches which have been shortened. Dotted lines represent long branches which have been shortened.

**Supplementary Figure 6.** A Ninja neighbour-joining tree showing the phylogenetic relationship between ST681 strains from this study and other publicly available strains that fell within the same HC1100 cluster (cgST complex). The study strains fell into two separate HC100 clusters, which are depicted in the two subtrees (B and C). The closest neighbours to both HC100 clusters are displayed, with the branch labels indicating the allelic distances between strains. The locations of the isolates are displayed for each tree, with Gambian strains highlighted in red. Dotted lines represent long branches which have been shortened.

**Supplementary Figure 7.** A phylogenetic tree showing the phylogenetic relationship between ST2076 strain (an MDR strain) and all other publicly available genomes that fell within the same HC1100 cluster (cgST complex). The legend shows the locations of the isolates, Gambian strains are highlighted in red. The subtree (B) shows the allelic distance between the study strain and its nearest relative, an ST2076 isolate recovered from water. Dotted lines indicate shortened long branches.

**Supplementary Figure 8.** A Ninja phylogenetic tree showing the closest neighbours of simian ST1873 strain—an environmental (soil) isolate belonging to ST83, separated from the study strain by 1659 alleles. The legends of both the main tree and the subtree show the locations of the isolates Gambian strains are highlighted in red. In the subtree (B), the closest neighbour to the simian ST1873 strain is also highlighted in red. Dotted lines are used to indicate shortened long branches.

**Supplementary Figure 9.** Ninja phylogenetic trees showing the closest neighbours to simian isolates belonging to novel sequence types (Achtman) ST8550 (A), ST8532 (B) and ST8525 (C), ST8826 (D). The allelic distances between these study isolates and their closest neighbours are >1100 alleles, and the closest neighbours belong to seven-allele STs which share less than five out of the seven MLST loci. Each genome (ST8550, ST8532, ST8525) belongs to a unique cgST complex (novel groups at HierCC 1100), indicative of novel diversity within the non-human primate niche.

## Supplementary files

**Supplementary File 1. A.** Characteristics of the study population, displaying the primate species, their age and gender, and the *E. coli* sequence types (Achtman MLST STs) and phylotypes recovered from individual samples. Novel STs are designated by an asterisk (*). **B.** Reference strains that were included in this study.

**Supplementary File 2. A.** Predicted plasmids from short-read sequences, using ARIBA PlasmidFinder (Reference 44). **B.** A table indicating the virulence and (or) resistance genes located on representative plasmids that were sequenced by Oxford nanopore technology. The size of each plasmid and the functions of the respective genes encoded thereon are also indicated.

**Supplementary File 3. A.** A summary of the sequencing statistics of the novel sequence types derived from this study. **B.** Prophage types detected from long-read sequences using PHASTER (reference 51).

**Supplementary File 4.** A summary of the sequencing statistics of the study isolates.

**Supplementary File 5.** List of virulence factors detected using ARIBA VFDB (Reference 44).

**Supplementary File 6.** Pair-wise single nucleotide polymorphism distances calculated from the core genome alignment using snp-dists v0.6 (https://github.com/tseemann/snp-dists).

