## Supplementary File 1A and B for "Genomic diversity of *Escherichia coli* isolates from non-human primates in the Gambia"

**Supplementary File 1A: Characteristics of the Study primate population**

| Sample ID | Gender | Age | Colony picks | Recovered STs (colonies per ST) |
| --- | --- | --- | --- | --- |
| PapRG-03 | F | Adult | 5 | ST336 (n=5) |
| PapRG-04 | M | Adult | 4 | ST1204 (n=2), ST8826* (n=1), ST1665 (n=1) |
| PapRG-05 | U | Juvenile | 4 | ST1431 (n=2), ST99 (n=1), ST6316 (n=1) |
| PapRG-06 | M | Adult | 5 | ST8827* (n=1), ST8525*, ST4080 (n=1), ST2521 (n=1), ST1204 (n=1) |
| ProbRG-07 | F | Adult | 5 | ST73 (n=5) |
| ChlosRG-12 | M | Adult | 4 | ST8824 (n=1), ST196 (n=2), ST40 (n=1) |
| ChlosAN-13 | F | Adult | 4 | ST8550* (n=1), ST8526* (n=1), ST1973 (n=2) |
| PapAN-14 | F | Adult | 5 | ST226 (n=3), ST2076 (n=1), ST939 (n=1) |
| PapAN-15 | F | Adult | 5 | ST8823 (n=1), ST5073 (n=1), ST226 (n=2), ST126 (n=1) |
| ChlosAN-17 | F | Juvenile | 4 | ST362 (n=1), ST681 (n=3) |
| ChlosAN-18 | F | Juvenile | 5 | ST681 (n=4), ST349 (n=1) |
| ProbAN-19 | F | Adult | 1 | ST8825* (n=1) |
| ChlosBP-21 | F | Adult | 5 | ST677 (n=4) |
| ChlosBP-23 | F | Adult | 3 | ST8527* (n=2), ST3306 (n=1) |
| ChlosBP-24 | M | Adult | 5 | ST73 (n=5) |
| ChlosBP-25 | U | Adult | 5 | ST3 (n=5) |
| ChlosM-29 | U | Adult | 2 | ST1873 (n=2) |
| PapM-31 | F | Adult | 5 | ST2800 (n=1), ST1727 (n=1), ST5780 (n=2), ST135 (1) |
| PapM-32 | F | Adult | 5 | ST8532* (n=1), ST212 (n=4) |
| PapM-33 | M | Adult | 5 | ST8533* (n=4), ST38 (n=1) |
| PapM-34 | M | Adult | 4 | ST676 (n=4) |
| PapM-36 | F | Adult | 2 | ST8535* (n=2) |
| PapKW-44 | U | Adult | 4 | ST442 (n=4) |
| ProbK-45 | F | Adult | 5 | ST127 (n=5) |
| Total |  |  | 101 |  |

Novel Sequence Types are indicated by an asterisk (*); M, Male; F, Female; U, Unknown.

**Supplementary File 1B: Reference strains used in this study**

| Strain | Sequence Type | Phylogroup designation | | GenBank assembly accession |
| --- | --- | --- | --- | --- |
| K-12 strain MG1655 | ST10 | | A | NC_000913.3 |
| 536 | ST127 | | B2 | GCA_000013305.1 |
| UMN026 | ST597 | | D | GCA_000026325.2 |
| IAI39 | ST62 | | F | GCA_000026345.1 |
| O157:H7 str. EDL933 | ST11 | | E | GCA_000732965.1 |
| IAI1 | ST1128 | | B1 | GCA_000026265.1 |
| IHE3034 | ST95 | | B2 | GCA_000025745.1 |
| *Escherichia fergusonii* | ST5298 | | Outroot species | GCA_000026225.1 |
